## Supplemental Materials for "Modeling hepatitis C micro-elimination among people who inject drugs with direct-acting antivirals in metropolitan Chicago"

**Table S1. HCV model infection parameters.**

| Description | Value | Range | Refs |
|---|---|---|---|
| Duration of acute infection in a naïve individual | 102 | 77-127 | (1, 2 ) |
| Duration of acute infection in a recovered individual | 28 | 8-48 | (2 ) |
| Time taken for an infected individual to become infectious | 3 | 2-4 | (2 ) |
| Probability of a recovered individual clearing virus upon re-exposure | 0.85 | 0.75-0.95 | (2, 3) |
| Probability of spontaneous viral clearance upon first exposure- females | 0.346 | 0.30-0.40 | (4) |
| Probability of spontaneous viral clearance upon first exposure - males | 0.121 | 0.10-0.14 | (4) |
| Probability that a naive PWID will be infected in a receptive sharing event with an infected PWID | 0.01 | 0.0005-0.05 | (5, 6) |

**Table S2. Parameters for the generation of the synthetic population.**

| Parameter description | Value | Range | Notes |
|---|---|---|---|
| Probability of chronic infection | 0.67 | 0.49-0.74 | (2, 7) |
| Attrition rate (per year) | 0.024 | 0.01-0.08 | (8) |
| Burn in days | 365 | - | Calibrated by observing the time necessary for the HCV incidence to stabilize. |
| Initial PWID population | 32,000 | 30,000-34,000 | (9) |
| Mean injection career duration (years) | 30.3 | 10-35 | (8) |
| Probability of cessation | 0.232 | 0.13-0.33 | (8) |
| Probability of acute HCV-infected PWID at time of inclusion | 0.05 | 0.01-0.09 | 0.9% prevalence among newer PWID, and 9% among other* PWID (10) |

*PWID who acquired HCV prior to initiating into injection drug use through other modes (e.g. sharing non-injection drug paraphernalia such as snorting straws.
